# Phytochemical Screening, Radical Scavenging and Antimicrobial Activities of the Crude Ethanolic, Methanolic, Ethyl Acetate and Chloroform Leaf Extracts of Arayat Pitogo (*Cycas riuminiana* Porte ex Regel)

**DOI:** 10.1101/2025.06.21.656729

**Authors:** Renato A. Dela Peña, Mohamed Afil Sabeer Hussain, Ashok Danaphal, Mohamed Daad Mohamed, Christian R. Pangilinan, Marilyn T. Malison, Angelico G. Reyes, Angelo B. Bañares, Monaliza M. Cayanan, Jacqueline V. Bagunu, Evelyn V. Totaan, Darwin E. Totaan, Daniel E. Gracilla

**Author notes:** Corresponding Author: Daniel Gracilla. Denotes Equal Contribution (Renato A. Dela Peña, Mohamed Afil Sabeer Hussain, Ashok Danaphal, Mohamed Daad Mohamed, Daniel E. Gracilla).

## Abstract

In the Philippines, *Cycas* spp. are found in Luzon Island particularly in Pampanga, Batangas, Bataan and Isabela provinces.. In this study, the bioactive potentials of the crude methanolic, ethanolic, ethyl acetate and chloroform leaf extracts of *Cycas riuminiana* Porte ex Regel were investigated. Based on the results of the four solvents used, the best extraction solvent for the phytochemicals is ethanol, followed by methanol, ethyl acetate and chloroform. The ethanolic and methanolic leaf extracts showed comparable antioxidant activity. The chloroform and ethyl acetate extracts also have comparable antioxidant activity but significantly lower than both methanolic and ethanolic extracts. However the greatest antimicrobial activity was exhibited by the ethyl acetate extract, followed by chloroform, methanolic and ethanolic extracts. The variation and similarity in the antioxidant and antimicrobial activities of the different extracts can be attributed to different mechanisms of interactions, namely, independent joint action, additive, synergistic, competitive or antagonistic interactions, among the bioactive compounds present in the crude extracts. Further studies are needed to elucidate the structure of the different phytochemicals present in the leaf extracts of Arayat Pitogo (*C. riuminiana* Porte ex Regel) and the specific mechanisms of interaction among these phytochemicals.

**SUMMARY:** The extracts were tested for the presence (trace, moderate or abundant amounts) of flavonoids, saponins, tannins, triterpenes, alkaloids, sterols and glycosides. The ethanolic extract was positive for all phytochemicals screened with sterols, flavonoids, glycosides and tannins being abundant, alkaloids being moderate and triterpenes and saponins in trace amounts. The methanolic extract was also positive for all constituents but in trace amounts, except for flavonoids which were abundant. The ethyl acetate extract contained abundant sterols, moderate alkaloids and trace amounts of saponins, glycosides and tannins. Finally, the chloroform extract contained abundant sterols, and trace amounts of alkaloids, saponins and glycosides.

The radical scavenging assay revealed that the highest percent inhibition was obtained for the ethanolic leaf extract (60.53±0.7801%), followed by methanolic extract (59.92±3.160%), chloroform extract (50.17±4.779%) and ethyl acetate extract (47.25±3.759%).

In terms of antibacterial activity, the ethyl acetate extract registered the highest inhibition against the three test organisms, namely, *Staphylococcus aureus, Bacillus subtilis* and *Escherichia coli*. The chloroform extract inhibited *S. aureus* and *B. subtilis*. The methanol extract inhibited *S. aureus* only. Finally, the ethanolic extract failed to inhibit any of the test organisms despite its abundant phytochemicals and high antioxidant activity.

In terms of antifungal activity, the different extracts inhibited *Candida albicans* with the ethyl acetate and chloroform extracts showing a high degree of inhibition followed by the methanolic and ethanolic extracts. However, none of the extracts showed any bioactivity against *Aspergillus niger*.

## INTRODUCTION

*Cycas*, the only genus under the family Cycadaceae, are found in many countries including the Philippines (1). Several species of *Cycas* were reported to thrive in the Philippines, namely, *Cycas saxatilis, C. aenigma, C. vespertilio, C. nitida, C. lacrimans, C. wadei, C. curranii, C. circinalis subsp. riuminiana var. curranii forma graminea* J. Schust. and *C. circinalis subsp. riuminiana var. curranii forma maritima* J. Schust (2). Some *Cycas*, like *C. circinalis*, have nut shells reported to possess medicinal property such as anti-bacterial activity against *Salmonella typhi* and *S. aureus* while the nut kernel was found effective against *Mycobacterium phlei* (3). Other *Cycas* can produce carcinogenic and neurotoxic compounds like cycasin from *C. revoluta* (4).

Many of these *Cycas* species have not been investigated for their medicinal potentials. One such species, *C. riuminiana* Porte ex Regel, has not been subjected to pharmacological potential and bioactivity analysis. This *Cycas* species is found in the provinces of Pampanga, Isabela, Bataan, and Batangas in Luzon Island of the Philippines (5). However, chemical profiling of the dichloromethane extract from the different parts of *C. riuminiana* yielded the following: from leaflets - α-tocopherol, phytyl fatty acid ester, squalene, lutein, chlorophyll a, long chain 1-alkene, and linoleic acid; from petiole, rachis and roots - β-sitosterol and stigmasterol; from sarcotesta – triacylglycerol; and from endotesta - β-sitosterol, triacylglycerol, methyl fatty acid ester and β-sitosteryl fatty acid ester (6).

Different solvents can extract different phytochemicals based on their polarity. The bioactivity of the crude extracts will depend upon the extracted phytochemicals working either synergistically or antagonistically (7). Thus, the use of different solvents may be considered in investigating the medical potentials of a plant. Indeed, the differential bioactive potentials of *C. riuminiana* Porte ex Regel may be revealed using different extraction solvents of varying polarities. Thus, four different solvents, namely, ethanol, methanol, ethyl acetate, and chloroform, were used to determine the phytochemical constituents and radical scavenging, antibacterial and antifungal activities of *C. riuminiana* Porte ex Regel.

## MATERIALS AND METHODS

### Research Design

To achieve the objective of this study, a series of laboratory procedures were carried out involving phytochemical extraction and screening and assays to determine the scavenging, antibacterial and antifungal activities of the extracts.

### Research Procedure

#### Collection and Plant Identification

The plant samples were collected from Mt. Arayat, in Magalang, Pampanga. The specimen was previously submitted for taxonomic authentication (8). Only leaves sample were taken from the plant and no plant was uprooted or died during the process of leaf collection.

#### Drying and Soaking

The leaves were air dried at room temperature for 7 days. The dried leaves were then grounded using a blender equipped with blade cutter. The ground leaves were divided into four parts and each part was soaked for two days in ethanol, methanol, ethyl acetate and chloroform in a 1:1 ratio.

#### Filtration and Rotary Evaporation

The soaked leaves were filtered, and the filtrates were then subjected to rotary evaporation at 40^°^C until all solvent was evaporated.

#### Phytochemical Screening

The phytochemical screening of the crude ethanolic, methanolic, chloroform and ethyl acetate extracts was conducted in the Industrial Technology Development Institute, Standards and Testing Division of the Department of Science and Technology, Philippines following the procedure according to Evans, Evans and Trease (2002) (9).

Extracts were screened for the presence of secondary metabolites. *Flavonoids* were detected using Mg^+^ turning test in which 1 mL of the extract was treated with 1.0 mL 10% HCl and Mg turnings. Red color formation indicates the presence of flavonoids. *Saponins* were determined by froth formation test in which the alcoholic extract was dissolved in hot water and then filtered. Vigorous shaking of the filtrate rendered it frothy whose honeycomb nature persisted for at least 30 minutes. For *tannin* presence, ferric chloride test was performed. Dried extract was dissolved in hot water and filtered. 1-2 drops of ferric chloride was added. Positive test is indicated by the appearance of dark coloration which may either be black, dark-blue or blue-black. *Triterpene* was detected by Liebermann-Burchard Test wherein a small amount of dried extract was dissolved in acetic anhydride followed by decantation of the soluble portion. 1-2 drops of concentrated sulfuric acid was later added. The formation of pink color turning to red indicates the presence of triterpenoids. Wagner’s test was carried out for *alkaloids* by dissolving a small amount of the dried extract in 1.0 mL of dilute acetic acid. The formation of cream precipitate indicates alkaloid presence. NaCl was added to remove false positive reactions. *Sterols* were tested by Salkowski’s Test in which concentrated sulfuric acid was added to 3 mg of the dried extract, and addition of chloroform with two drops of acetic acid. Blue color formation indicates the presence of sterols. To screen for *glycosides*, the Fehling’s Test was performed. The alcoholic extract was first dissolved in hot water and then filtered. 2 mL of the filtrate was placed in two test tubes one of which served as the control. 1 mL dilute HCl was added to one test tube but none in the control test tube. The two test tubes were placed in a boiling water bath for 5 minutes. Upon cooling, the mixtures were neutralized using anhydrous sodium carbonate until effervescence stopped. 1 mL of Fehling’s solution was added to the test tubes which were then heated in water bath for 2 minutes. An increased amount of brick red precipitate in the hydrolyzed test tube indicates the presence of glycoside.

#### Radical Scavenging Activity

The DDPH Free Radical Scavenging Assay performed in this study was adapted from Molyneux (2004) (10). A 300 µM free – radical solution was prepared by dissolving 1 mg of 2,2-diphenyl-1-(2,4,6-trinitrophenyl) hydrazyl (DDPH) in 10 ml Absolute ethanol. The extract prepared at 95 µl was dispensed to 96-well microtiter plates. Gallic Acid was used as positive control while DMSO was used as negative control. Five microliters of the controls and test samples were added to the wells to make a final volume of 100 µl. The plate was then incubated in dark and ambient temperature for 60 minutes. After incubation, absorbance was read at 570 nm. Based on the absorbance readings, the free radical inhibition of the test sample was computed using the formula: % Inhibition = [(Absorbance (-) control – Absorbance Sample) / (Absorbance (-) control – Absorbance gallic acid)] *100

#### Antimicrobial Sensitivity Test

Antibacterial Testing and Antifungal Testing were done as previously described (17), The bacterial test organisms, namely, *Bacillus subtilis* ATCC 6633, E*scherichia coli* and *Staphylococcus aureus* were obtained from the Department of Public Health Medical Microbiology, University of the Philippines, Manila. The *S. aureus* and *E. coli* strains used in the study were clinical isolates previously identified by biochemical analyses. The fungal test organisms, namely, *Candida albicans* UPCC 2168 and *Aspergillus niger* UPCC 4219 were provided by the Natural Science Research Institute, University of the Philippines, Diliman, Quezon City. A loopful of each test organism was inoculated into 2.5 ml sterile peptone water contained in 5cm test-tube (10mm diameter) and compared to 0.5 McFarland standard `to` give an approximately 1.0 × 10^8^ cfu/ml of the test bacterium and fungus. DMSO at 0.5% concentration was used as solvent control (SC). Tetracycline (coded as T+) with a quantity of 30 µL (concentration: 10µg/ml) per well was set as positive control for antibacterial activity while Canesten solution was used as positive control for antifungal activity. Double distilled water was used as negative control (coded as T0).

### Treatment of Data

Statistical analyses were conducted using GraphPad Prism ver. 6.1. To test for the presence of significant differences among the leaf extracts in terms of free radical scavenging activity, antibacterial and antifungal activity, one-way Analysis of Variance with Tukey’s Multiple Comparison Test as post-hoc test was conducted.

## RESULTS

### Phytochemicals Present in the Four Crude Leaf Extracts of Arayat Pitogo (*C. riuminiana*)

Results show (Table 1) that the ethanolic extract has abundant amounts of sterols, flavonoids, glycosides and tannins, moderate amounts of alkaloids and trace amounts of triterpenes and saponins. The methanolic leaf extract has abundant amount of flavonoids and trace amounts of sterols, triterpenes, alkaloids, saponins, glycosides and tannins. The chloroform leaf extract has abundant amount of sterols. Trace amounts of alkaloids, saponins and glycosides are present, but triterpenes and tannins were absent. The ethyl acetate leaf extract has abundant amount of sterols, moderate amount of alkaloids, trace amounts of saponins, glycosides and tannins. The different extracts contained variable amounts of the phytochemicals screened.

**TABLE 1.** Phytochemical Test Results for the Various Extracts

| Phytochemicals<br>Screened | Solvent Used for Extraction |  |  |  |
| --- | --- | --- | --- | --- |
|  | Ethanol | Methanol | Chloroform | Acetate |
| Sterols | (+++) | (+) | (+++) | (+++) |
| Triterpenes | (+) | (+) | (-) | (-) |
| Flavonoids | (+++) | (+++) | (-) | (-) |
| Alkaloids | (++) | (+) | (+) | (++) |
| Saponins | (+) | (+) | (+) | (+) |
| Glycosides | (+++) | (+) | (+) | (+) |
| Tannins | (+++) | (+) | (-) | (+) |

Legend: (+) Trace, (++) moderate, (+++) abundant, (-) Absent

### Radical Scavenging Activity of the Four Crude Leaf Extracts of Arayat Pitogo (*C. riuminiana*)

Figure 1 and Table 2 show the percent of inhibition of each extract based on the DPPH free radical scavenging assay. Gallic acid was used as the positive control while DMSO as the negative control. The free radical scavenging activities of the crude extracts relative to gallic acid were assayed using the 1,1-diphenyl -2-picryl hydrazyl. The assay was performed three times (replicates) for each crude extract. As shown in Figure 1, the antioxidant activities of the four crude extracts are statistically lower than Gallic Acid (*p*<0.0001) but significantly higher than DMSO (*p*<0.0001). Multiple comparisons reveal that the antioxidant activities of the ethanolic extract is comparable with that of the methanolic extract (*p*=0.9998). The same holds true between ethyl acetate and chloroform extracts (*p*=0.7943). However, the antioxidant activities of both the ethanolic and methanolic extracts are significantly higher than both chloroform and ethyl acetate extracts (*p*<0.05). Thus, in terms of the radical scavenging activity, ethanolic extract = methanolic extract > chloroform extract = ethyl acetate extract. The results suggest that the varying antioxidant activities of the extracts may be due to the varying groups and concentrations of phytochemicals present in them.

**TABLE 2.** Computed Percent Inhibition of DPPH by the Different Extracts

| Extract | Trial 1 | Trial 2 | Trial 3 | Mean±SD |
| --- | --- | --- | --- | --- |
| Methanolic Extract | 60.21% | 56.62% | 62.92% | 59.92±3.160% |
| Ethanolic Extract | 61.38% | 60.35% | 59.85% | 60.53±0.7801% |
| Ethyl Acetate Extract | 49.71% | 42.92% | 49.11% | 47.25±3.759% |
| Chloroform Extract | 52.53% | 44.67% | 53.31% | 50.17±4.779% |

**Figure 1.**
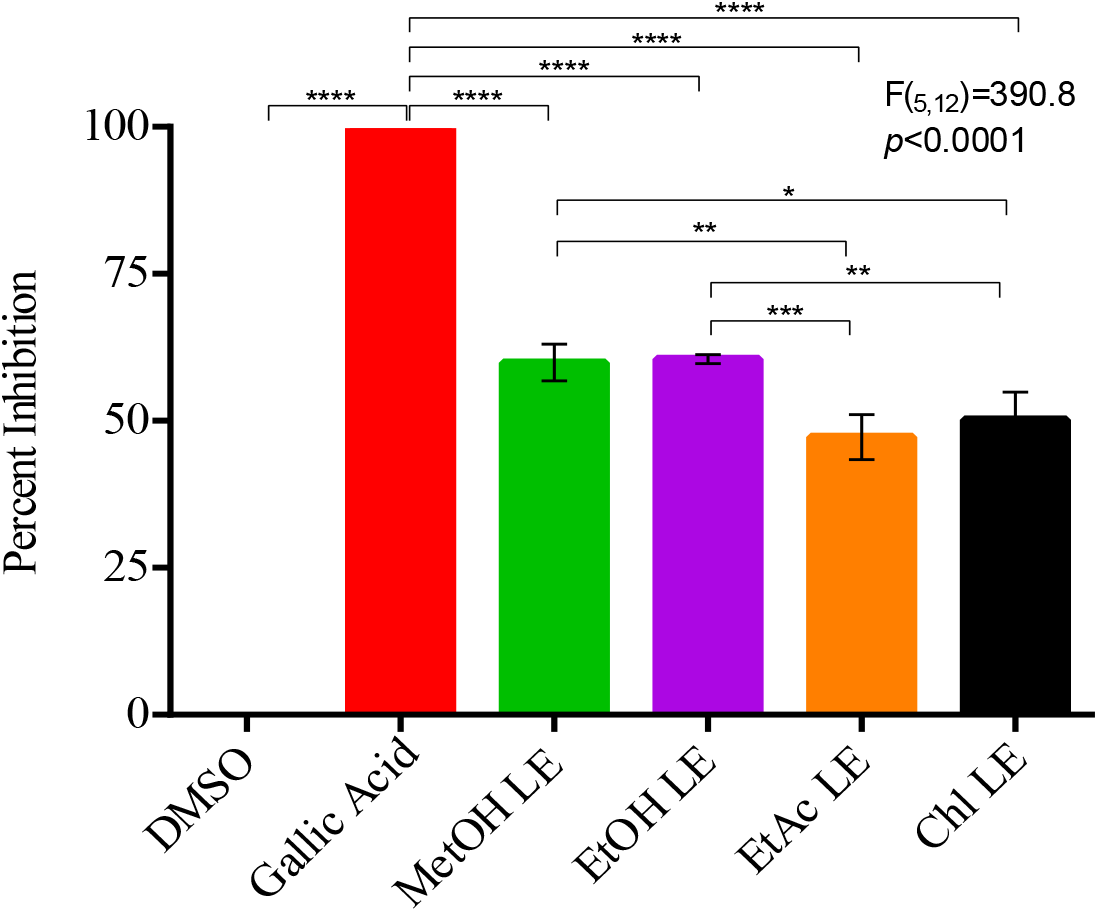
Comparison of the Radical Scavenging Activity of the Different Leaf Extractsxsxs

### Antibacterial Activity of Ethanolic, Ethyl Acetate, Methanolic and Chloroform Extracts

#### Inhibition of *E. coli* using Arayat Pitogo (*C. riuminiana* Porte ex Regel) crude leaf extracts

Shown in Figure 2 and Table 3 are the results of the antibacterial sensitivity testing of the different extracts against *E. coli*. Only T2 (ethyl acetate extract) can inhibit *E. coli* with an inhibitory activity comparable with the T+ (tetracycline) with inhibition zone diameters of 30.7 mm and 36.3 mm, respectively. The ability of T2 (ethyl acetate extract) to kill *E. coli* indicates that ethyl acetate is effective in extracting phytochemicals that can inhibit or suppress the growth of Gram-negative bacteria. This result also suggests that the antimicrobial compound/s against *E. coli* is/are polar.

**Table 3.** Zones of inhibition of the Different Extracts against *E. coli*

| Treatments | Mean Zone of Inhibition (mm)<sup>*</sup> |
| --- | --- |
| T- (Distilled Water) | 0<sup>a</sup> |
| T1 (Ethanol Extract) | 0<sup>a</sup> |
| T2 (Ethyl acetate Extract) | 30.7<sup>b</sup> |
| T3 (Methanol Extract) | 0<sup>a</sup> |
| T4 (Chloroform Extract) | 0<sup>a</sup> |
| T+ (Tetracycline) | 36.3<sup>c</sup> |
| Solvent Control (DMSO) | 0<sup>a</sup> |

<sup>\*</sup> Values with similar subscripts are not statistically different.

**Figure 2.**
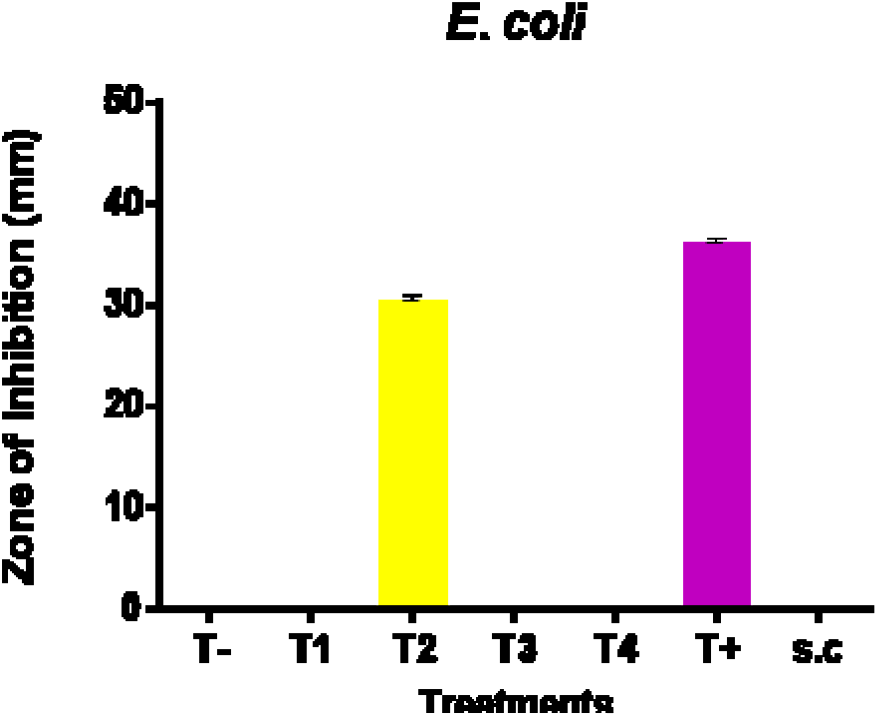
Comparison of the Means of Zone of Inhibition against *E. coli*

#### Inhibition of *S. aureus* using Arayat Pitogo (*C. riuminiana* Porte ex Regel) crude leaf extracts

Shown in Figure 3 and Table 4 are the results of the antibacterial testing of the different extracts against *S. aureus*. T2 (ethyl acetate extract), T3 (methanol extract), and T4 (chloroform extract) can inhibit *S. aureus* with zone of inhibition diameters of 32.7 mm, 28 mm, 15.3 mm and 35.5 mm, respectively. T+ (tetracycline) is still significantly more effective than the crude extracts with T2>T3>T4. The inhibitory action of T2 (ethyl acetate extract), T3 (methanol extract) and T4 (chloroform extract) against *S. aureus* indicates the presence of various polar and nonpolar antibacterial compounds which possibly exert an independent joint action, additive and/or synergistic action against Gram-positive bacteria.

**Table 4.** Zones of Inhibition of Different Extracts against *S. aureus*

| Treatments | Mean Zone of Inhibition (mm)<sup>*</sup> |
| --- | --- |
| T- (Distilled Water) | 0<sup>a</sup> |
| T1 (Ethanolic Extract) | 0<sup>a</sup> |
| T2 (Ethyl Acetate Extract) | 32.7<sup>d</sup> |
| T3 (Methanolic Extract) | 28<sup>c</sup> |
| T4 (Chloroform Extract) | 15.3<sup>b</sup> |
| T+ (Tetracycline) | 35.5<sup>e</sup> |
| Solvent Control (DMSO) | 0<sup>a</sup> |

\*Values with similar superscripts are not statistically different.

**Figure 3.**
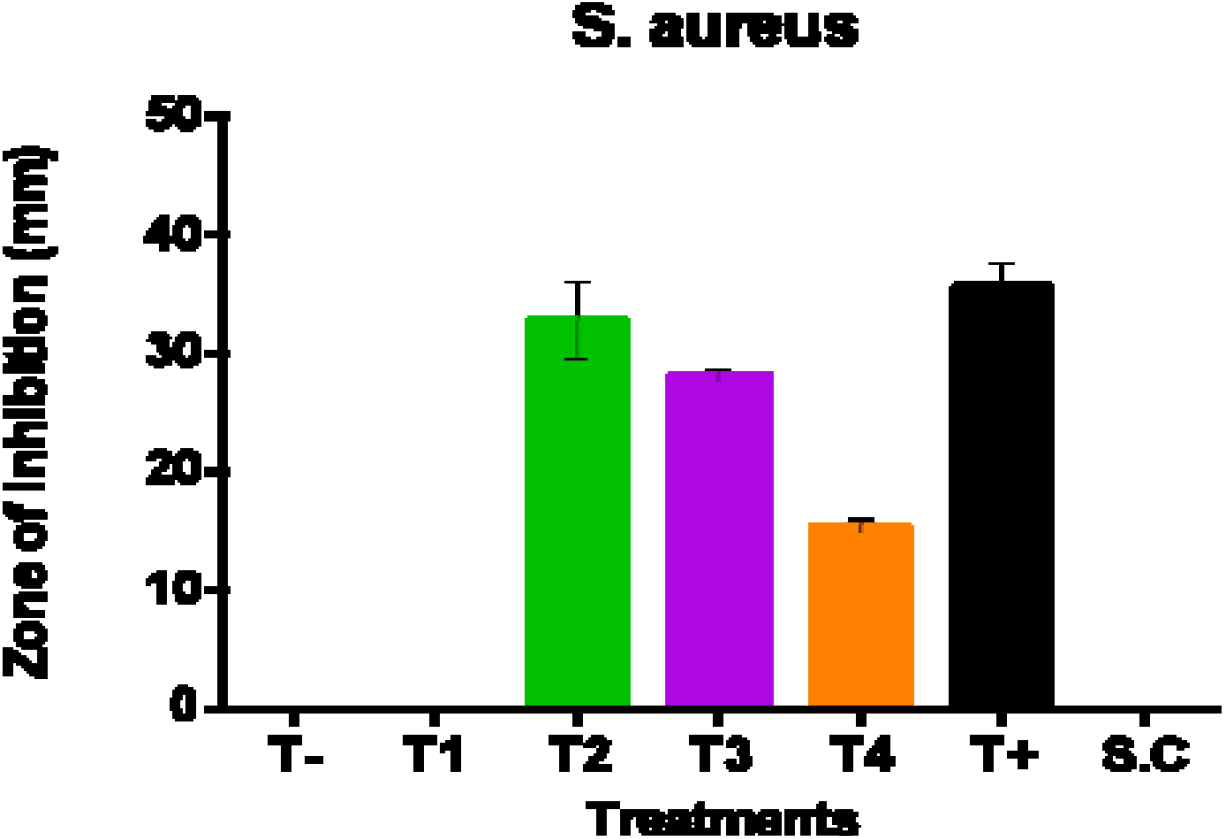
Comparison of Means of Zones of Inhibition against *S. aureus*

#### Inhibition of *B. subtilis* using Arayat Pitogo (*C. riuminiana* Porte ex Regel) crude leaf extracts

Shown in Figure 4 and Table 5 are the results of the antibacterial testing of the different extracts against *B. subtilis*. Of the four extracts, only T2 (ethyl acetate extract) and T4 (chloroform extract) inhibited *B. subtilis* with zone of inhibition diameters of 30 mm and 18 mm, respectively. T+ had a zone of inhibition diameter of 34.3 mm. The inhibition of *B. subtilis* by T2 (ethyl acetate extract) and T4 (chloroform extract) indicates the presence of various phytochemicals with differing polarities that are bioactive against Gram-positive, spore-forming bacteria.

**Table 5.**
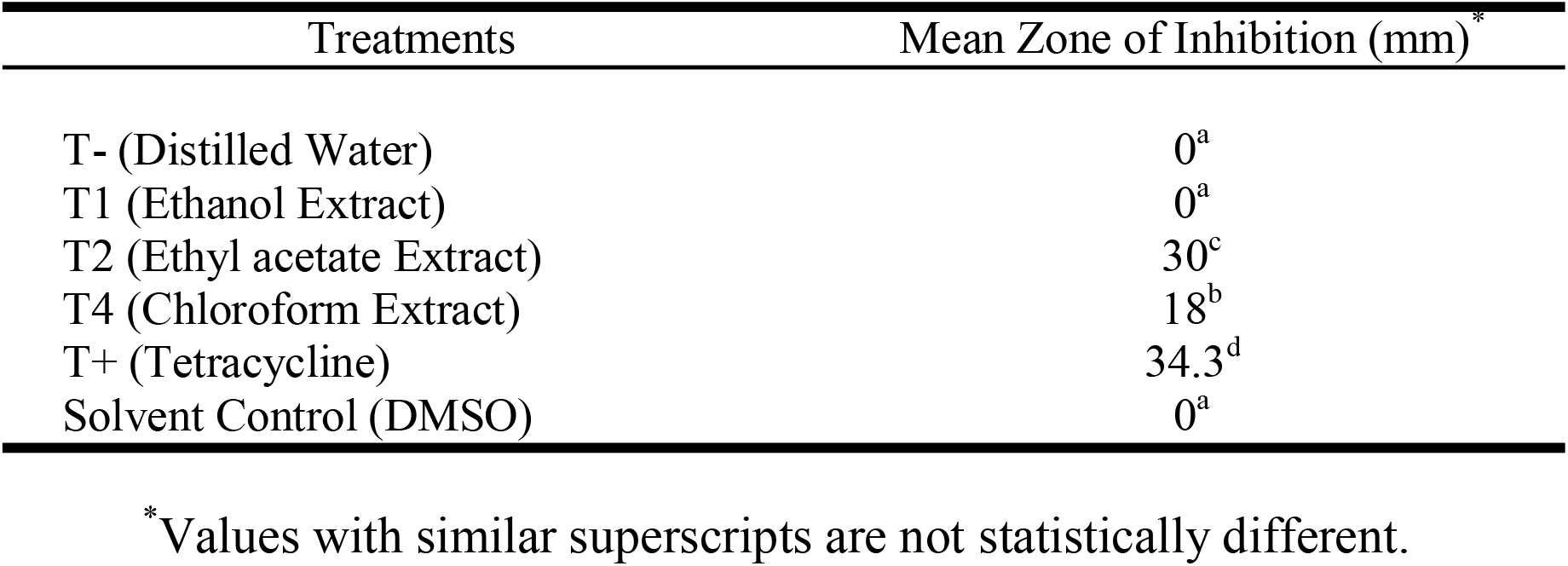
Zones of Inhibition of Different Extracts against *B. subtilis*

| Treatments | Mean Zone of Inhibition (mm)* |
| --- | --- |
| T- (Distilled Water) | 0<sup>a</sup> |
| T1 (Ethanol Extract) | 0<sup>a</sup> |
| T2 (Ethyl acetate Extract) | 30<sup>c</sup> |
| T4 (Chloroform Extract) | 18<sup>b</sup> |
| T+ (Tetracycline) | 34.3<sup>d</sup> |
| Solvent Control (DMSO) | 0<sup>a</sup> |

\*Values with similar superscripts are not statistically different.

**Figure 4.**
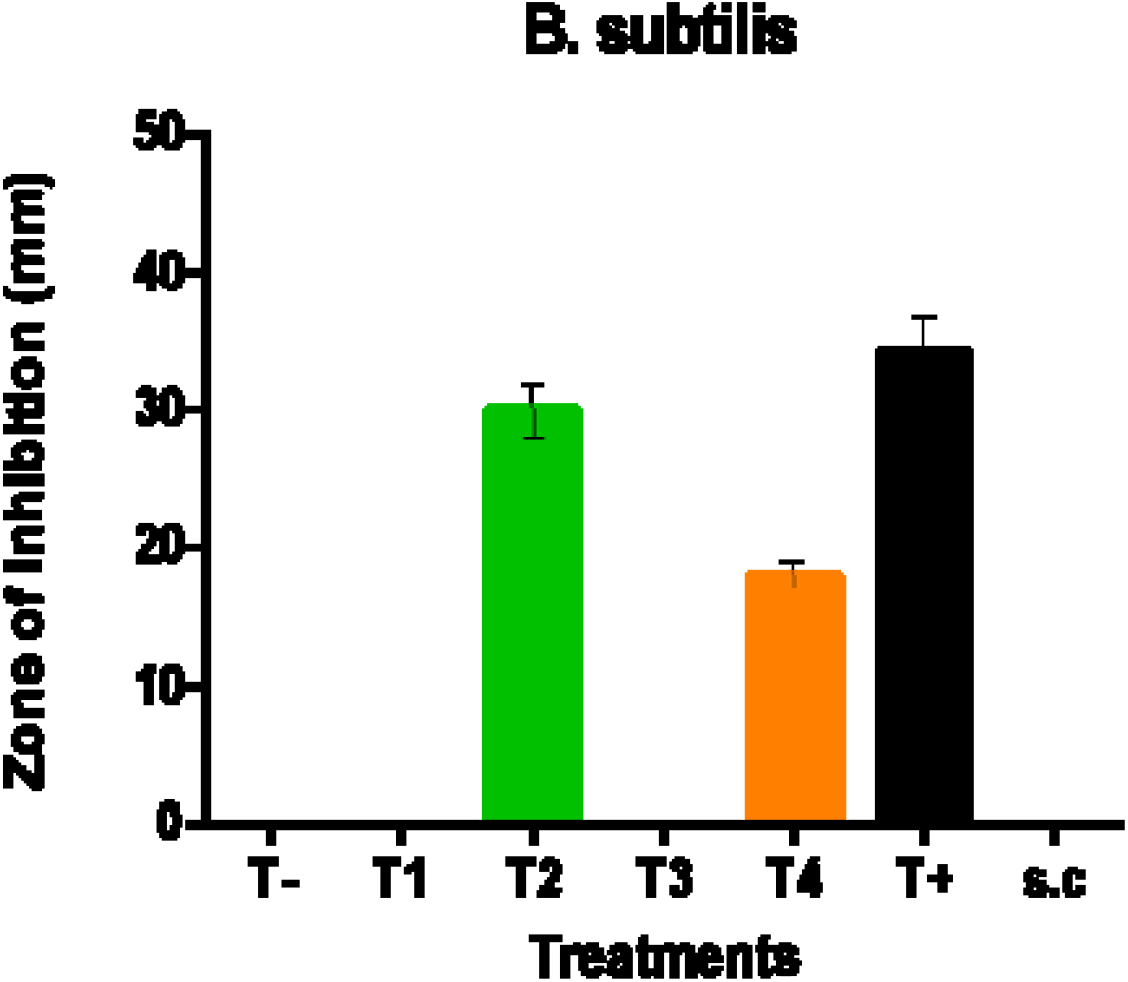
Comparison of Means of Zones of Inhibition against *B. subtilis*

### Antifungal Activity of Ethanolic, Ethyl Acetate, Methanolic and Chloroform Extracts

The antifungal activity of the four extracts were tested against *C. albicans* and *A. niger*. As shown in Figure 5 and Table 6, all four extracts have comparable inhibitory potential against *C. albicans*. However, T+ (Canesten solution) was significantly more effective than the four extracts. Finally, none of the extracts inhibited *A. niger*. These results indicate that there are antifungal compounds in Arayat Pitogo with possibly varying polarities.

**Table 6.** Zone of inhibition of different extract against *Candida albicans*

| Treatments | Mean Zone of Inhibition (mm)* |
| --- | --- |
| T- (Distilled water) | 0<sup>a</sup> |
| T1 (Ethanol Extract) | 12.8<sup>b</sup> |
| T2 (Ethyl acetate Extract) | 13.9<sup>b</sup> |
| T3 (Methanol Extract) | 13.0<sup>b</sup> |
| T4 (Chloroform Extract) | 13.9<sup>b</sup> |
| T+ (Tetracycline) | 32.0<sup>c</sup> |
| Solvent Control (DMSO) | 0<sup>a</sup> |

\*Values with similar superscripts are not statistically different.

**Figure 5.**
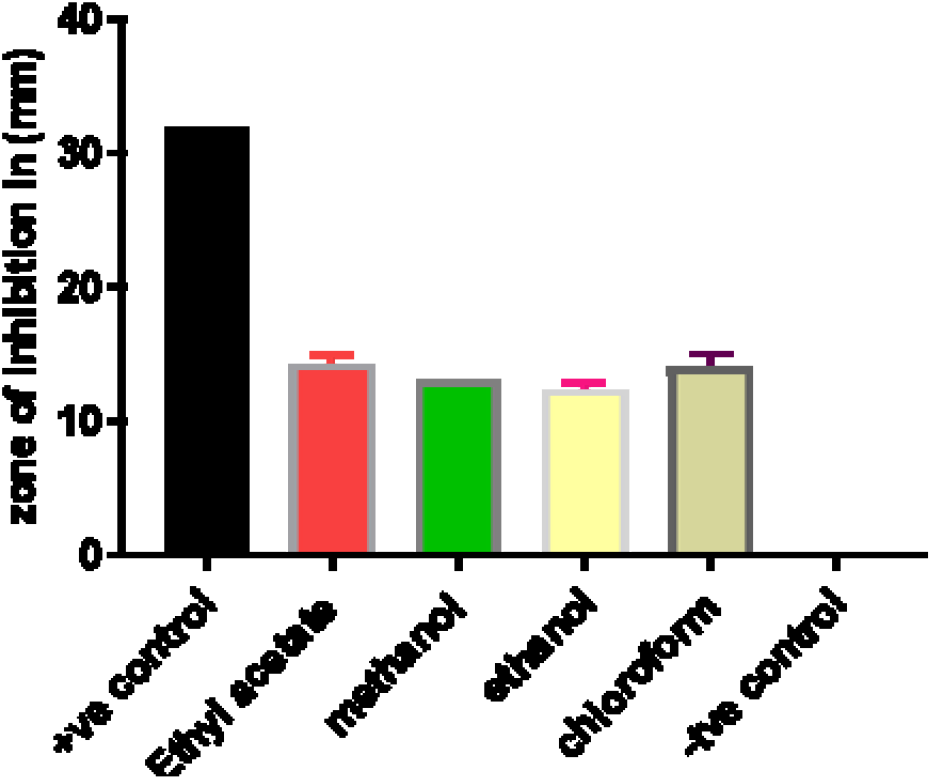
Comparison of Antifungal Activity of Different Extracts of *Cycas riuminiana* against *Candida albicans*

### Summary of Bioactivities of the Phytochemicals in the Different Extracts of *C. riuminiana*

Table 7 shows that the sterols, alkaloids, saponins, glycosides and tannins in the ethyl acetate extract appear to have inhibitory action against *S. aureus, B. subtilis, E. coli* and *C. albicans* but are ineffective against *A. niger*. Tannins appear to be necessary for bioactivity against *E. coli* since its absence in the chloroform extract rendered the extract inactive against *E. coli*. The ethyl acetate and chloroform extracts differed only in tannins. The relatively lower antioxidant activity of the two extracts is probably due to the absence of flavonoids and triterpenes. The presence of triterpenes (trace) and flavonoids (abundant) in both the methanolic and ethanolic extracts appear to have substantially reduced the bioactivity of these extracts against *B. subtilis* and *E. coli* by possibly interfering with the activity of the other phytochemicals, particularly tannins. The tannins of the methanolic extract are present in trace amount while the tannins in the ethanolic extract are present in abundant amount. The presence of triterpenes and flavonoids probably inactivated the tannins. However, the triterpenes and flavonoids account for the significantly higher antioxidant activity of these two extracts. The bioactivity of all the extracts against *C. albicans* may be due to the independent joint action, additive or synergistic interactions among the sterols, alkaloids, saponins, and glycosides which are commonly present in all the extracts. However, none of these phytochemicals have any bioactivity against *A. niger* possibly because of competitive or antagonistic interactions among them.

**Table 7.** Summary of Bioactivities and Phytochemical Constituents of the Different Crude Leaf Extracts of Arayat Pitogo (*Cycas riuminiana* Porte ex Regel)

| Parameters Studied | LEAF EXTRACTS OF ARAYAT PITOGO |  |  |  |
| --- | --- | --- | --- | --- |
| <sup>1</sup>Bioactivity |  |  |  |  |
| 1. Antibacterial Activity | Ethyl Acetate | Chloroform | Methanolic | Ethanolic |
| *S. aureus* | + | + | + | - |
| *B. subtilis* | + | + | - | - |
| *E. coli* | + | - | - | - |
| 2. Antifungal Activity |  |  |  |  |
| *C. albicans* | + | + | + | + |
| *A.Niger* | - | - | - | - |
| 3. Antioxidant Activity | 47.25±3.76% | 50.17±4.78% | 59.92±3.16% | 60.53±78% |

<sup>2</sup>Phytochemicals Screened

|  |  |  |  |  |
| --- | --- | --- | --- | --- |
| Sterols | (+++) | (+++) | (+) | (+++) |
| Triterpenes | (-) | (-) | (+) | (+) |
| Flavonoids | (-) | (-) | (+++) | (+++) |
| Alkaloids | (++) | (+) | (+) | (++) |
| Saponins | (+) | (+) | (+) | (+) |
| Glycosides | (+) | (+) | (+) | (+++) |
| Tannins | (+) | (-) | (+) | (+++) |

Legend: <sup>1</sup>For Bioactivity: + (bioactive), - (inactive)
<sup>2</sup>For Phytochemicals: (+) Trace, (++) moderate, (+++) abundant, (-) Absent

## DISCUSSION

In solvent-based extraction, the variability of the phytochemical constituents may depend on the polarity of the extraction solvents, the polarity of the metabolites to be extracted (11) and rigidity of the plant sample. Based on dielectric constant, both methanol and ethanol are semi-polar solvents while ethyl acetate and chloroform are considered nonpolar solvents (12). Methanol has been reported to be the solvent that can best extract most of the phytochemicals including alkaloid, flavonoid, saponins and tannins (13). Ethanol, which is commonly used for extraction, can also extract various phytochemicals (13). Ethyl acetate was considered to be the best extraction solvent for tannin, but it can also extract alkaloid, flavonoids and other phytochemicals while chloroform is preferred for non-polar compounds (13). Our results show that in extracting phytochemicals from Arayat Pitogo (*C. riuminiana*), ethanol extracted all the target phytochemicals with moderate to abundant amounts except for triterpenes and saponins. Methanol can extract all the tested phytochemicals but in lesser or trace amounts except for flavonoids (extracted in abundant amount). Methanol (CH_3_OH) and ethanol (CH_3_CH_2_OH), which are both polar protic solvents with a polar group (-OH) and non-polar tail, are capable of dissolving compounds with polar protic structure but methanol is more polar than ethanol (12). The phytochemicals they dissolve are more polar in nature. Ethyl acetate and chloroform, on the other hand, are nonpolar solvents as they contain no -OH group but chloroform is specifically a chlorinated solvent (12). Since nonpolar solvents dissolve nonpolar solutes, the phytochemicals present in the ethyl acetate and chloroform extracts are more nonpolar.

In terms of the target phytochemicals in *C. riuminiana*, our results show that the sterols can be extracted in abundant amounts using either ethyl acetate, chloroform, or ethanol while methanol would yield trace amounts only. Triterpenes can be extracted in trace amount using either ethanol or methanol. Flavonoids can be extracted in abundant amounts using either ethanol or methanol. Alkaloids can be extracted in moderate amounts using either ethanol or ethyl acetate and in trace amounts using either chloroform or methanol. Saponins can be extracted in trace amounts using any of the four solvents. Glycosides can be extracted in abundant amounts using ethanol but in trace amounts using any of the other three solvents. Finally, tannins can be extracted in abundant amounts using ethanol but in trace amounts using either ethyl acetate or methanol. These results can be used as a preliminary guide by future researches who may intend to concentrate on using specific solvents to extract specific phytochemicals. We have not attempted to use different combinations of the solvents in this study.

Based on our results, the high abundance of flavonoids in the ethanolic and methanolic extracts of *C. riuminiana* may principally account for these extracts’ higher radical scavenging potential compared to chloroform and ethyl acetate extracts. Sterols appear to be most abundant in ethyl acetate and chloroform extracts and, together with the other phytochemicals in trace amounts, may principally account for the radical scavenging activity of these extracts. Most flavonoids, tannins, triterpenes, saponins, and glycosides have antimicrobial properties (15). Many alkaloids (15) and many plant sterols (16) also have antimicrobial properties. However, the complex interaction among the bioactive compounds in the form of “independent joint action, additive effects, antagonistic effects and synergistic effects” (14) may affect the antimicrobial action of the crude extracts. These are at work in the four crude extracts of *C. riuminiana* in our study. The inhibitory potential of the ethyl acetate extract is consistently higher against *E. coli, S. aureus*, and *B. subtilis* compared to the other extracts. It is also the only extract that inhibited *E. coli*. This inhibitory action of the ethyl acetate extract may be attributed to the independent joint action, additive or synergistic interactions among the sterols (abundant), alkaloids (moderate), saponins (trace), glycosides (trace) and tannins (trace). The chloroform extract, with phytochemical composition almost similar to ethyl acetate extract except for alkaloids in trace amount, is effective against *S. aureus* and *B. subtilis* but inactive against *E. coli*. The methanolic extract is effective only against *S. aureus*. Notably, it contains abundant flavonoids but trace amounts of sterols, triterpenes, alkaloids, saponins, glycosides and tannins. The ethanolic extract, while containing the most abundant amounts of phytochemicals, did not inhibit any of the bacterial test organisms. All extracts inhibited *C. albicans* but not *A. niger*. These results suggest that competitive or antagonistic interactions are also occurring among the phytochemicals in the extracts, particularly, in the ethanolic extract.

The varying interactions among the phytochemicals in the extracts of *C. riuminiana* may be due to bioavailability, interference with cellular transport processes, activation of pro-drugs or deactivation of active compounds to inactive metabolites, action of synergistic partners at different points of the same signalling cascade (multi-target effects) or inhibition of binding to target proteins (7). The present study, however, was not aimed to specifically identify which mechanisms are at work among the phytochemicals in the extracts. But based on the results of the antimicrobial activity, the phytochemicals in the ethyl acetate extract of *C. riuminiana* are most likely working either through independent joint action, additive or synergistic interactions while those in the ethanol extract are most likely interacting either competitively or antagonistically. For the chloroform and methanolic extracts, these possible mechanisms may also be at work but at varying levels or degrees. These results suggest that the antimicrobial compounds present in *C. riuminian*a could be differentially extracted using different solvents, or combination of solvents, of varying polarities.

## CONCLUSION

The different phytochemicals extracted from the leaves of Arayat Pitogo (*C. riuminiana* Porte ex Regel) using different solvents of varying polarities and the complex interactions among these phytochemicals may account for the observed similarity and variation in the radical scavenging and antimicrobial activities of the ethyl acetate, chloroform, methanolic and ethanolic crude extracts. The results suggest that the best extraction solvent for the phytochemicals from the leaves of Arayat Pitogo (*C. riuminiana* Porte ex Regel) is ethanol, followed by methanol, ethyl acetate and chloroform. The antioxidant activities of the ethanolic and methanolic extracts were comparable to each other but significantly higher than both ethyl acetate and chloroform extracts due to the presence of more phytochemicals. The different extracts, however, showed variable antibacterial activity against *S. aureus* (ethyl acetate, methanol and chloroform extracts), *B. subtilis* (ethyl acetate and chloroform extracts), and *E. coli* (ethyl acetate extract only) but similar antifungal activity against *C. albicans* (all crude extracts) and *A. niger* (all negative), which can be attributed to different interactions among the bioactive compounds present in the crude extracts. The results suggest that independent joint action, synergistic or additive interactions may be occurring primarily in the ethyl acetate extract, which exhibited the greatest antimicrobial activities, while competitive or antagonistic interactions operate mainly in the ethanol extract, which showed the least antimicrobial activity. The various mechanisms of interactions among the phytochemicals may be occurring in varying degrees in the chloroform and methanolic extracts. These results call for follow-up studies, especially, on the ethyl acetate extract since it exhibited the highest antimicrobial activities. Further studies are needed for the structural elucidation of the bioactive compounds present in the crude leaf extracts of Arayat Pitogo (*C. riuminiana* Porte ex Regel) and to elucidate the mechanism of interactions among these phytochemicals in the extracts.

The aim of this study is to inform the public about the promising pharmaceutical and antimicrobial potential of the `Cycas` *riuminiana Porte ex Regel*. The study did not further elucidate the MIC concentration as we utilized different solvent to guide the other researcher which solvent is the most effective for extracting antimicrobial compound which ethyl acetate. do

## Abbreviations

NCBI: (National Center for Biotechnology Information)
BLAST: (Basic Local Alignment Search Tool)
16S rRNA gene: (16 Svedberg ribosomal ribonucleic acid)
cfu: (colony forming unit)
TNTC: (too numerous to count)
NA: (Nutrient Agar)
MHA: (Muller Hinton Agar)
HSD: (honest significant difference)
ANOVA: (Analysis of Variance).

## Acknowledgement

Acknowledgement is due to the Office of Research and Development, Manila Central University, UPD Natural Science Research Institute and ITDI, Standards and Testing Division, DOST-Taguig.

